# Criticality-related dynamics accompany long-term functional decline in primary cortical MEA cultures

**DOI:** 10.64898/2026.10.09.757871

**Authors:** Ane Escobar, Francisco Javier Gil-Bea, Laura Roderiguez-Gómez

## Abstract

Primary cortical neuron cultures are widely used to study network development and decline, but their long-term electrophysiological trajectories are poorly characterized. We used multi-electrode arrays to longitudinally monitor Wistar rat cortical cultures for up to ∼730 days in vitro (DIV), combining activity measures with survival analysis and criticality metrics. Wells shared an early developmental phase, with mean firing rate peaking at 12–14 Hz around DIV 100–150, before diverging into two phenotypes: early-death wells, showing progressive loss of burst organization, longer inter-burst intervals, shorter bursts, and steep declines in active-electrode recruitment; and long-survival wells, which sustained organized bursting and measurable activity well beyond DIV 400, in some cases past DIV 700. Cox proportional-hazards modeling with FDR correction across thirteen candidate features identified burst duration, inter-burst interval, synchrony, mean firing rate, branching ratio, and Kappa index as significant predictors of time to network failure. Criticality metrics offered the most mechanistic insight: in early-death wells, branching ratio declined toward zero and Kappa index rose toward its upper bound as networks approached silencing. Beyond their average levels, the rate of change of branching ratio and Kappa index over recordings was the strongest correlate of survival time among all criticality metrics. Neuronal avalanche size and duration exponents and their scaling relation, remained consistent with scale-free dynamics in both phenotypes while networks were active, though avalanche occurrence rate declined progressively in early-death wells without reaching significance as an independent predictor. These results show that long-term network survival is not determined by firing rate alone, but by preserved burst organization, sustained electrode recruitment and near-critical propagation dynamics. This work offers one of the longest electrophysiological characterizations of rodent cortical cultures to date and identifies criticality-based metrics as promising tools for predicting network decline and evaluating interventions to preserve long-term neuronal function.

**Author summary:** Cultures of neurons grown on multi-electrode arrays let researchers record the electrical activity of developing brain-like networks over time. We recorded electrical activity from rat cortical neuron cultures continuously for up to nearly two years, one of the longest such studies to date. We found that networks followed one of two distinct trajectories: about a third of the networks lost their organized firing patterns within the first year in culture, while the rest sustained coordinated activity well beyond that point, in some cases for close to two years. Firing rate alone (how often neurons fired) did not reliably distinguish these two groups. What did was the structure of that activity: how firing was organized into bursts, how synchronized different regions of the network remained, and how efficiently electrical signals propagated across the network, a property linked to networks operating near a so-called “critical” state, thought to support efficient information processing in the brain. These features changed months before other, more obvious signs of decline appeared, in some cases within the first three to ten months in culture. We propose that tracking this kind of network-level organization, rather than raw activity levels, offers a more sensitive way to study neuronal ageing and to test whether experimental treatments genuinely protect brain network function.

## Introduction

Primary cortical neurons dissociated from embryonic rat brain and cultured on multi-electrode arrays (MEAs) have become one of the most versatile tools in in vitro neuroscience, enabling non-invasive, continuous and long-term recording of network electrophysiology^1,2^. These cultures self-organize into functional networks within days of plating, progressing through stereotyped developmental stages that closely mirror early in vivo cortical maturation^3,4^. The developmental trajectory of spontaneous firing in primary cortical cultures is well established over the first four to eight weeks in vitro. Following plating, neurons begin to extend neurites and form synaptic contacts, with the first action potentials typically appearing around days in vitro (DIV) 4–5^5^. During the second week, isolated spikes give way to coordinated bursts of activity and by DIV 14–21 most cultures exhibit mature synchronized bursting (SB) patterns^6,7^. The mean firing rate (MFR) rises sharply during this period, peaks and then stabilizes to a characteristic plateau^3^. Network synchrony, measured by indices such as the synchrony index (SI), also increases in parallel with bursting maturation^8,9^. Burst count per electrode follows a broadly similar trajectory, often peaking during the transition to mature network states before becoming more variable with time^10^.

Beyond the first month, relatively little is known about the long-term evolution of firing dynamics in primary rodent cortical cultures. Most published studies report recordings up to DIV 30–60^9,11^ and only a handful have sustained cultures on MEAs for several months^12,13^. Dichter et al. demonstrated that dissociated cortical neurons from embryonic rats can be maintained in vitro for 8–12 weeks while retaining mature electrophysiological properties^14^. More recently, the development of sealed culture chambers compatible with MEAs has enabled recordings exceeding one year^12^. Nonetheless, a detailed characterization of the temporal evolution of MFR, synchrony and burst dynamics across hundreds of DIV and across distinct preparations, has not been reported.

The reduced availability of long-term electrophysiological data is particularly limiting in the context of neuronal ageing research. In vitro neuronal cultures have been proposed as accessible models of neurodegeneration and aging^15,16^, yet their utility depends critically on understanding how network activity evolves not just during maturation, but across the full lifespan of the culture. While long-term neuroglial co-cultures have been validated as ageing models based on senescence markers and microRNA profiles^15(p)^, the electrophysiological correlates of this progressive decline and whether they can serve as functional biomarkers of network ageing, remain largely unexplored. This is an important gap, since MEA-based functional readouts offer a direct, non-invasive window into network health that biochemical assays cannot provide^17^.

From a translational perspective, the ability to track network survival over months or years in vitro opens the possibility of using these cultures as pharmacological ageing models for testing compounds that may delay or prevent neuronal decline. MEA platforms have already demonstrated utility for drug screening in the context of neurotoxicity^18^, epilepsy^19^ and neurodegenerative disease^20,21^. However, the use of electrophysiological network dynamics as a readout of ageing has not been systematically investigated. A model that characterizes the natural trajectory of network decline, identifies electrophysiological functional survival associated-features of survival and can detect compounds that shift wells from a short-survival to a long-survival phenotype, would represent a significant advance for the field.

Here we present a longitudinal characterization of primary rat individual cortical networks maintained on 24-well MEA plates for up to two years in vitro. We describe the full temporal evolution of firing rate, burst dynamics, synchrony, electrode recruitment and criticality metrics and identify electrophysiological signatures that characterize electrophysiological trajectories associated with different functional-survival times. In addition to conventional activity measures, we examine branching ratio, Kappa index, DFA exponent, the coefficient of variation of the inter-spike interval, Fano factor and neuronal avalanche statistics, including avalanche occurrence rate and avalanche size/duration exponents, to determine whether near-critical network organization provides a more sensitive window into long-term network fate. We further show that specific parameters, in particular burst organization, active electrode retention, criticality metrics during early and intermediate recording windows and the longitudinal trajectory (slope) of criticality metrics, correlate significantly with long-term survival. Cox proportional-hazards modelling with false-discovery-rate correction provides significant univariate time-to-event-based confirmation for a subset of these functional survival associated-features and we establish the practical limits of avalanche-exponent fitting once avalanche counts decline, suggesting that MEA-based electrophysiology can serve as a predictive biomarker of network ageing in vitro. Taken together, our findings provide a quantitative framework for using long-term cortical MEA cultures as a model of neuronal ageing and as a platform for electrophysiology-guided screening of neuroprotective or anti-ageing compounds.

## Results

### Well viability and survival classification

A total of 48 wells were monitored across two independent 24-well MEA plates. After applying the prespecified viability filter, defined as at least 8 of 16 electrodes with a firing rate greater than 1 Hz during DIV 20–40, 34 wells were retained for analysis and 14 were excluded. Based on the predefined functional network-failure criterion, the 34 retained wells were retrospectively classified according to whether they met this criterion before or after DIV 365: 13 were classified as early-death wells and 21 as long-survival wells (**Table S1**). Plate 1 contributed 9 early-death and 11 long-survival wells, whereas Plate 2 contributed 4 early-death and 10 long-survival wells.

For descriptive visualization, Kaplan–Meier curves were generated for these two operationally defined groups (**Figure 1A**). As expected from the classification criterion, all early-death wells met the functional network-failure criterion before DIV 365, whereas long-survival wells met it later or remained functionally active until the end of the study.

**Figure 1.**
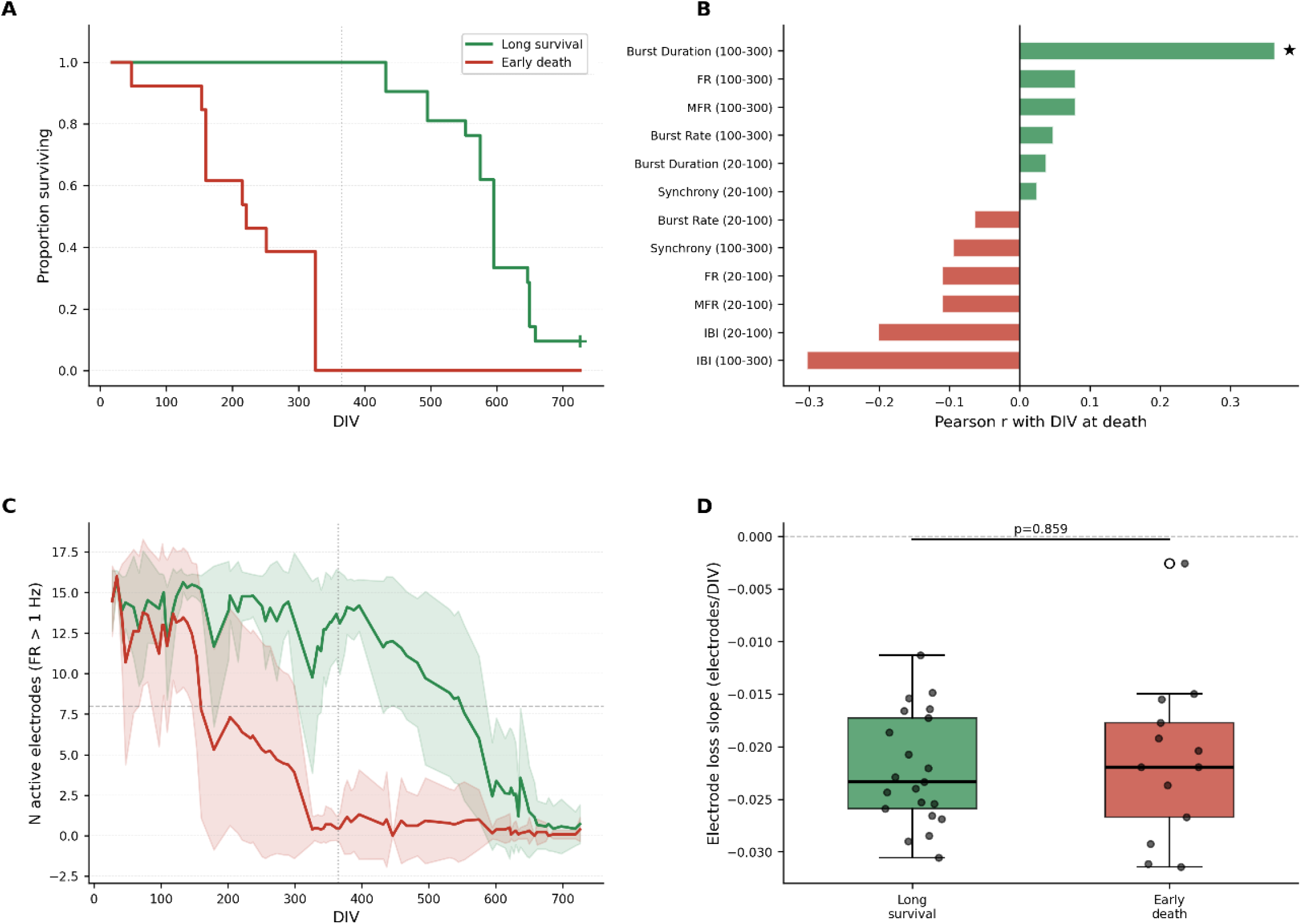
Survival analysis and electrophysiological metrics associated with neuronal network longevity. (A) Kaplan–Meier survival curves showing the proportion of surviving neuronal cultures over time. Wells were classified as long survival (green) or early death (red) according to the predefined survival criterion based on the day in vitro (DIV) at which the network met the death criterion. The vertical dotted line indicates DIV 365 (1 year in vitro), and the tick mark on the green survival curve denotes censored observations corresponding to cultures that remained viable until the end of the study. (B) Pearson correlation coefficients between electrophysiological metrics and the DIV at which each neuronal culture reached the predefined death criterion. Correlations were calculated for three temporal windows (DIV 20–100, DIV 20–300, and DIV 100–300), as indicated for each metric. Positive and negative correlation coefficients are displayed separately, and statistically significant correlations (P < 0.05) are indicated by an asterisk. (C) Temporal evolution of the number of active electrodes per well (firing rate > 1 Hz) in neuronal cultures classified as long survival (green) or early death (red). Solid lines represent the group mean at each day in vitro (DIV), and shaded regions indicate ±1 standard deviation (SD). The horizontal dashed line indicates the activity threshold used for well inclusion (≥8 active electrodes), and the vertical dotted line marks DIV 365 (1 year in vitro). (D) Distribution of the electrode loss rate for individual wells, calculated as the slope of the linear regression between the number of active electrodes and time (electrodes/DIV). Each point represents one well. Boxes indicate the interquartile range (IQR), the central line represents the median, and whiskers extend to 1.5 × IQR. The p-value for the comparison between long-survival and early-death groups is shown above the boxplots.

For descriptive visualization, Kaplan–Meier curves were generated for these two operationally defined groups (**Figure 1A**). As expected from the classification criterion, all early-death wells met the functional network-failure criterion before DIV 365, whereas long-survival wells met it later or remained functionally active until the end of the study. Two long-survival wells were therefore censored at the end of follow-up. For all longitudinal analyses reported, each well’s contribution to the group-level trajectories ends at the DIV on which that well met the death criterion (or at censoring, for the long-survival wells still active at study end); no post-death or post-censoring activity is included, so any divergence between the two groups reflects genuine differences in network dynamics while both phenotypes were still electrophysiologically active.

### Functional survival associated-features and electrode loss

To identify early electrophysiological signatures of later network fate, we computed Pearson correlations between each metric averaged in two early windows (DIV 20–100 and DIV 100–300) and the DIV at which each well reached the death criterion; results appear in **Figure 1B**. The strongest positive association with survival was burst duration in the DIV 100–300 window (r ≈ 0.38, p < 0.05), indicating that wells displaying longer bursts in months 3–10 tended to survive longer. In contrast, firing-rate metrics (including mean firing rate, MFR) were comparatively weak functional survival associated-features in the earlier windows (r ≈ −0.10 to −0.12). Inter-burst interval (IBI) was the clearest negative functional survival associated-feature (r ≈ −0.15 to −0.30, most negative at DIV 100–300); higher IBI (sparser bursting) in the early windows associated with shorter survival, suggesting that early loss of compact burst structure forecasts premature network failure.

To validate these associations with a method that explicitly accounts for censoring and the timing of network death, we fit univariate Cox proportional-hazards models for thirteen candidate features (burst, activity and criticality metrics averaged within DIV 20–100 and/or DIV 100–300) and corrected for multiple comparisons using the Benjamini–Hochberg procedure (**Table 1**; **Table S4**). Seven features reached FDR-corrected significance (q < 0.05): Kappa index (100–300, HR = 3.02), branching ratio (100–300, HR = 0.34), inter-burst interval (100–300, HR = 1.38), mean firing rate (20–100, HR = 1.36), synchrony (20–100, HR = 0.74), burst duration (100–300, HR = 0.78) and synchrony (100–300, HR = 0.87). Notably, mean firing rate during DIV 20–100, which showed only a weak simple correlation with survival time (**Figure 1B**), emerged as a significant Cox functional survival associated-feature (higher early firing rate associated with earlier death), indicating that the survival-model framework is sensitive to modest but consistent early differences that pairwise correlation alone does not resolve. Mean firing rate at DIV 100–300, burst rate (either window), burst duration at DIV 20–100, inter-burst interval at DIV 20–100 and avalanche occurrence rate at DIV 100–300 did not reach significance in this analysis (**Table 1**).

**Table 1.** Individual predictors of network death, combining Cox proportional-hazards results (FDR-corrected) with Pearson-correlation analyses.

| Metric | Category | Window (DIV) | Statistic | Direction of association | Significance | Source |
| --- | --- | --- | --- | --- | --- | --- |
| **Kappa index ($\kappa$)** | Criticality | 100–300 | Cox HR = 3.02 [2.41, 3.79] | Higher $\kappa \rightarrow$ earlier death | $q < 0.001$ | Table S4 |
| **Branching ratio ($\sigma$)** | Criticality | 100–300 | Cox HR = 0.34 [0.26, 0.44] | Lower $\sigma \rightarrow$ earlier death | $q < 0.001$ | Table S4 |
| **Inter-burst interval (IBI)** | Burst structure | 100–300 | Cox HR = 1.38 [1.27, 1.50] | Longer IBI → earlier death | q < 0.001 | Table S4 |
| **Mean firing rate (MFR)** | Activity | 20–100 | Cox HR = 1.36 [1.22, 1.51] | Higher early MFR → earlier death | q < 0.001 | Table S4 |
| **Synchrony** | Activity | 20–100 | Cox HR = 0.74 [0.67, 0.80] | Lower early synchrony → earlier death | q < 0.001 | Table S4 |
| **Burst duration** | Burst structure | 100–300 | Cox HR = 0.78 [0.71, 0.86] | Shorter bursts → earlier death | q < 0.001 | Table S4 |
| **Synchrony** | Activity | 100–300 | Cox HR = 0.87 [0.81, 0.93] | Lower synchrony → earlier death | q < 0.001 | Table S4 |
| **Branching ratio (slope)** | Criticality | Full recording | Pearson r = 0.48 | Declining $\sigma$ over time → earlier death | p < 0.05 | Fig. 3I |
| **CV of ISI (slope)** | Criticality | Full recording | Pearson r = 0.46 | n/a | p < 0.05 | Fig. 3I |
| **Kappa index (slope)** | Criticality | Full recording | Pearson r = -0.39 | Rising $\kappa$ over time → earlier death | p < 0.05 | Fig. 3I |
| **Burst duration** | Burst structure | 100–300 | Pearson r = 0.38 | Shorter bursts → earlier death | p < 0.05 | Fig. 1B |
| **Branching ratio ($\sigma$)** | Criticality | 100–300 | Pearson r = 0.28 | Lower $\sigma$ → earlier death | n.s. (r only) | Fig. 3I |
| **Kappa index ($\kappa$)** | Criticality | 100–300 | Pearson r = -0.27 | Higher $\kappa$ → earlier death | n.s. (r only) | Fig. 3I |
| **Mean firing rate (MFR)** | Activity | 100–300 | Cox HR = 1.22 [0.92, 1.64] | n/a | q = 0.279 (n.s.) | Table S4 |
| **Avalanche occurrence rate** | Criticality | 100–300 | Cox HR = 0.78 [0.52, 1.17] | n/a | q = 0.339 (n.s.) | Table S4 |
| **Inter-burst interval (IBI)** | Burst structure | 20–100 | Cox HR = 1.23 [0.75, 2.01] | n/a | q = 0.529 (n.s.) | Table S4 |
| **Burst rate** | Activity | 20–100 / 100–300 | Cox HR = 1.05 / 1.01 | n/a | n.s. | Table S4 |
| **Burst duration** | Burst structure | 20–100 | Cox HR = 1.03 [0.90, 1.17] | n/a | q = 0.748 (n.s.) | Table S4 |

Figure 1C shows the temporal evolution of active-electrode counts (FR > 1 Hz) in each group: long-survival wells retained a larger number of active electrodes for longer, whereas early-death wells exhibited an earlier and more rapid reduction in electrode recruitment. Individual electrode-loss slopes (Figure 1D) trended toward steeper declines in early-death wells, although the between-group comparison did not reach statistical significance in this data set.

### Longitudinal network activity

Figure 2 shows the longitudinal evolution of six electrophysiological activity metrics **(Table S2**) in cultures classified as long survival and early death. For both groups, the first 100–150 DIV were characterized by a common developmental trajectory: MFR increased from early baseline values, reached a peak around DIV 100–150 and then began to stabilize. This indicates that both phenotypes reached mature network activity states before their later divergence.

**Figure 2.**
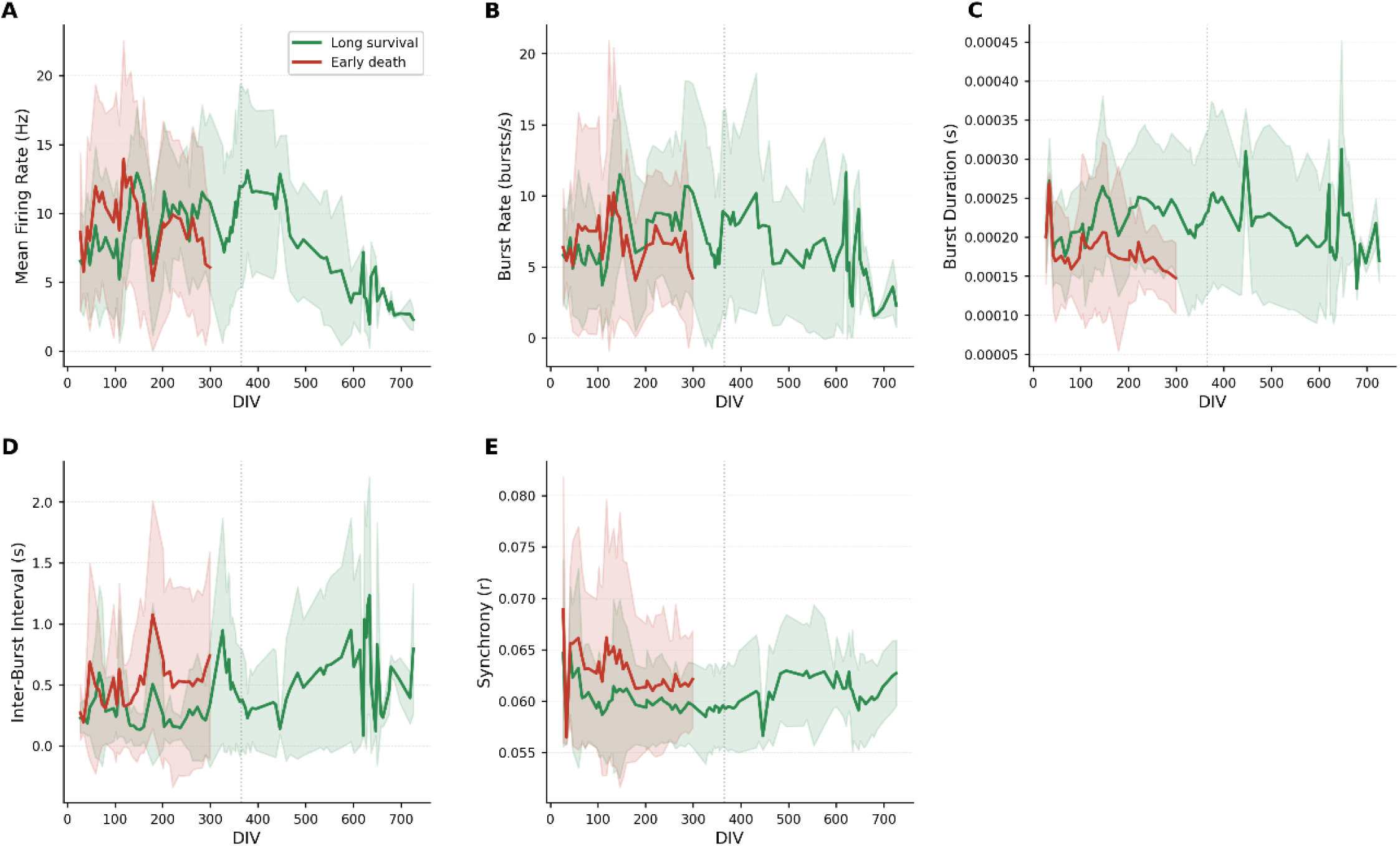
Longitudinal evolution of electrophysiological activity metrics in neuronal cultures. Longitudinal changes in electrophysiological activity metrics calculated from extracellular recordings of neuronal cultures classified as long survival (green) or early death (red). Solid lines represent the group mean at each day in vitro (DIV), and shaded regions indicate ±1 standard deviation (SD). The vertical dotted line marks DIV 365 (1 year in vitro). (A) Mean firing rate (MFR), calculated as the average firing rate across all active electrodes within each well. (B) Burst rate, expressed as the number of detected bursts per second. (C) Mean burst duration. (D) Mean inter-burst interval (IBI). (E) Synchrony, quantified as the mean pairwise Pearson correlation coefficient among electrodes within each well.

After this shared developmental phase, the trajectories separated. In long-survival wells, MFR remained measurable for much longer, with values fluctuating through the mid-to-late recording period before declining gradually toward the end of the experiment (Figure 2A). In early-death wells, MFR decreased earlier and more steadily, reaching low values much sooner. Burst rate followed a similar pattern (Figure 2B): both groups showed active bursting during maturation, but long-survival wells maintained burst activity for longer, whereas early-death wells showed an earlier collapse in burst frequency.

Burst duration also differed between groups (Figure 2C). Long-survival wells generally showed longer and more sustained bursts across the recording period, while early-death wells exhibited shorter burst durations and a clearer decline over time. Inter-burst interval was particularly informative (Figure 2D): early-death wells tended to show longer intervals between bursts, indicating less frequent and less stable burst organization. By contrast, long-survival wells maintained more compact burst timing for much longer.

Synchrony showed a comparatively smaller separation between groups (Figure 2E). Both phenotypes exhibited similar values during early development, but long-survival wells maintained slightly more stable synchrony later in the recording, whereas early-death wells showed a modest decline or flattening of this metric. Overall, Figure 2 indicates that the two groups share an initial maturation phase but diverge later in terms of burst organization, persistence of activity and long-term maintenance of network dynamics.

### Criticality analysis: distinct dynamical regimes between survival groups

Figure 3 presents longitudinal criticality metrics (**Table S3**), computed from recordings obtained after DIV 18, together with windowed and correlation-based comparisons between groups. Dashed blue reference lines in each panel mark the reference values commonly used in the corresponding analytical framework for each metric (σ = 1, κ = 1, H = 0.5 and Fano factor = 1), against which the group trajectories can be interpreted.

**Figure 3.**
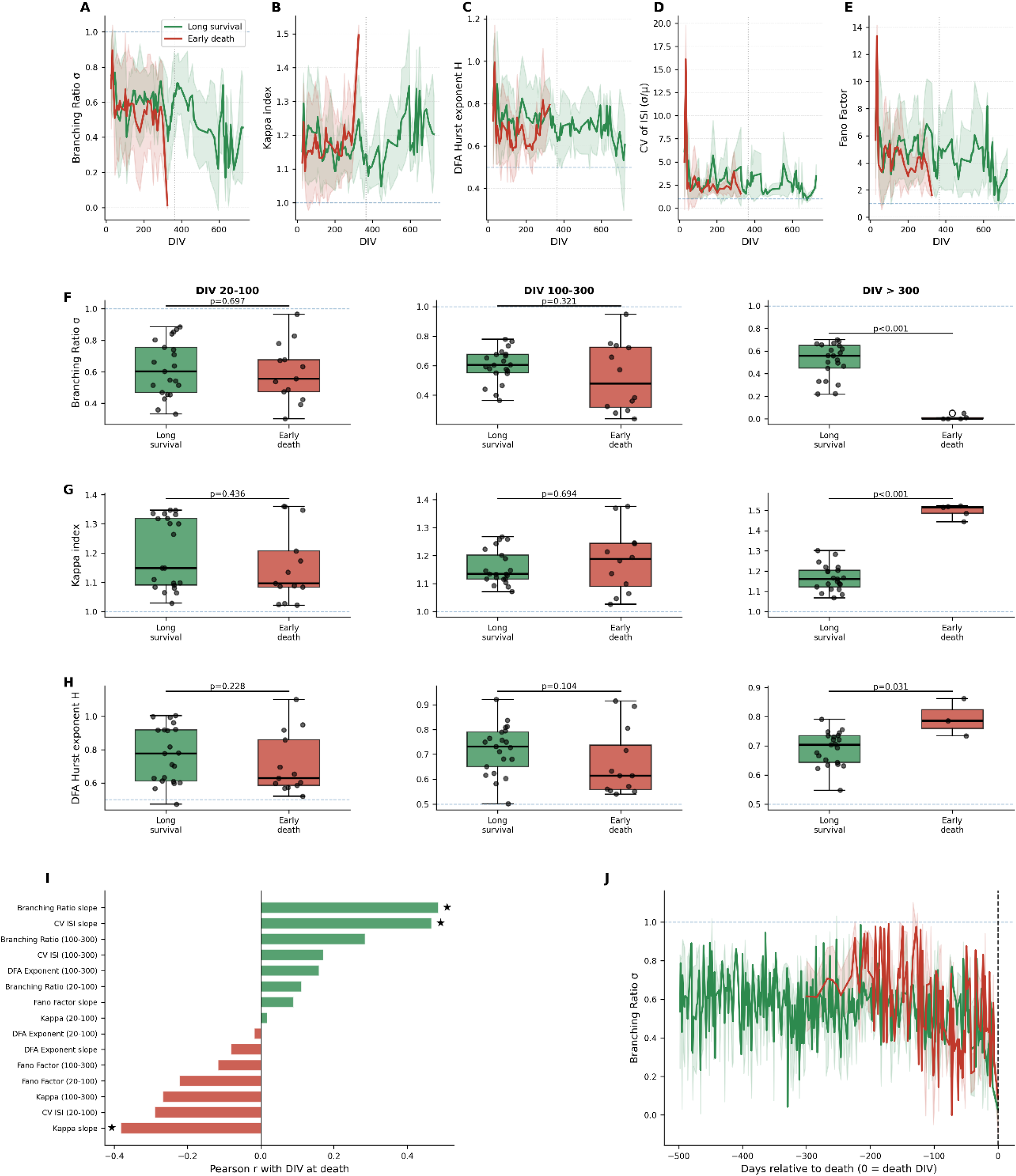
Temporal evolution of criticality metrics in long-surviving and early-death neuronal cultures. Mean ± standard deviation (SD) of the principal criticality metrics calculated from extracellular spike trains of neuronal cultures classified as long survival (green) or early death (red). Analyses include only recordings obtained after DIV 18. (A) Branching ratio (σ). (B) Kappa index (κ). (C) Detrended fluctuation analysis (DFA) exponent (H). (D) Coefficient of variation of interspike intervals. (E) Fano factor (variance/mean of spike counts). Solid lines represent the group mean at each day in vitro (DIV), and the shaded regions indicate ±1 SD. Horizontal dashed blue lines denote the theoretical critical values for each metric (σ = 1, κ = 1, H = 0.5, and Fano factor = 1). The vertical dotted line marks DIV 365 (1 year in vitro). (F, G, H) Columns correspond to the temporal windows used for the analysis (DIV 20–100, DIV 100–300, and DIV >300), while rows display the branching ratio (σ), kappa index (κ) and detrended fluctuation analysis (DFA) exponent (H), respectively. Each point represents an individual well. Boxes indicate the interquartile range (IQR), the central line represents the median, and whiskers extend to 1.5 × IQR. Horizontal dashed blue lines indicate the theoretical critical values for each metric (σ = 1, κ = 1, and H = 0.5). Statistical comparisons between groups were performed independently for each time window, and the corresponding P values are shown above each comparison. (I) Pearson correlation coefficients between criticality-derived metrics and the day in vitro (DIV) at which each neuronal culture reached the predefined death criterion. Correlations were calculated for multiple temporal windows preceding culture death (DIV 20–100, 20–300, and 100–300, as indicated). Positive and negative correlation coefficients are displayed separately, and statistically significant correlations (P < 0.05) are indicated by an asterisk. (J) Temporal evolution of the branching ratio (σ) aligned to the day of culture death (defined as day 0). Individual trajectories are shown for cultures classified as long survival (green) and early death (red), with the dashed vertical line indicating the reference death time. The x-axis represents days relative to culture death, allowing comparison of branching ratio dynamics preceding network failure independently of the absolute DIV.

The branching ratio σ (Figure 3A) diverges between groups: both start subcritical (∼0.5–0.7 at DIV 20–50) and early-death wells decline toward zero by the time they reach their individual death DIV (∼DIV 300–350), while long-survival wells maintain intermediate σ values below the critical line for much longer, declining only gradually across the full ∼730-DIV recording. The Kappa index κ (Figure 3B) shows the reciprocal pattern: both groups start near mildly supercritical values (∼1.1–1.2), but early-death wells rise toward the ceiling (∼1.5) as they approach their own death DIV, consistent with loss of scale-free avalanche structure and emergence of large, disorganized events, whereas long-survival wells remain closer to the κ = 1 critical line for most of the recording, with only a modest late rise.

DFA-derived Hurst exponent H (Figure 3C) overlaps between groups for most of the recording (H ∼0.6–0.8, above the H = 0.5 critical line in both), indicating preserved long-range correlations; H rises further in early-death wells in the period immediately preceding death, indicating a shift toward increasingly non-stationary, less scale-free dynamics as these networks approach failure. The coefficient of variation of the inter-spike interval (Figure 3D) provides a complementary measure of spike-timing irregularity: it is markedly elevated in early-death wells during the first ∼50 DIV (transiently exceeding 15) before settling into a range that overlaps with long-survival wells for most of the remaining recording. The Fano factor (Figure 3E) follows a broadly similar pattern, starting high in both groups and settling to FF ∼2–8, with long-survival wells tending toward the higher end of this range later in the recording, reflecting greater spike-count variability in networks that remain active for longer.

Windowed comparisons (Figure 3F**–H**) test these differences directly across three temporal windows (DIV 20–100, DIV 100–300 and DIV >300) for branching ratio, Kappa index and DFA exponent, respectively. Group differences were not statistically significant in either the DIV 20–100 window (σ: p = 0.697; κ: p = 0.436; H: p = 0.228) or the DIV 100–300 window (σ: p = 0.321; κ: p = 0.694; H: p = 0.104), when both phenotypes were still broadly overlapping. Differences became significant only in the DIV >300 window (σ: p < 0.001; κ: p < 0.001; H: p = 0.031), which is populated predominantly by long-survival wells together with the small number of early-death wells that survived long enough to contribute data close to their own death; in this window, early-death wells show branching ratio collapsed near zero and Kappa saturated near its 1.5 ceiling, consistent with terminal network collapse rather than early prediction.

In the correlation analysis (Figure 3I), the strongest individual criticality-based functional survival associated-features of survival time were the longitudinal slopes of branching ratio (r ≈ 0.48, p < 0.05) and of the CV of the inter-spike interval (r ≈ 0.46, p < 0.05), together with the slope of the Kappa index (r ≈ −0.39, p < 0.05). No window-averaged criticality metric (branching ratio or Kappa at DIV 20–100 or DIV 100–300, DFA exponent, or Fano factor) reached significance by simple correlation, although branching ratio (100–300, r ≈ 0.28) and Kappa (100–300, r ≈ −0.27) showed the same directional trend and were also supported in univariate Cox analysis as significant functional survival associated-features by the Cox proportional-hazards model (**Table 1**). This indicates that, in addition to their absolute level within a fixed window, the trajectory of these metrics over time, whether branching ratio and CV of ISI are rising or falling and whether Kappa is drifting toward its ceiling, carries independent functional survival associated-feature.

Finally, branching ratio trajectories aligned to the day of death (Figure 3J) show that, in the months preceding death, early-death wells transiently reach higher, at times near-critical or mildly supercritical, branching-ratio values than long-survival wells at the equivalent aligned offset, before undergoing an abrupt collapse toward zero in the final weeks before network silencing. Long-survival wells fluctuate around a comparatively lower and noisier, but more stable, subcritical range across the same aligned window, with a more gradual decline as they approach their own (typically much later) death or censoring point.

### Avalanche size and duration statistics

Neuronal avalanches, cascades of threshold-crossing spikes bounded by silent periods, provide a finer-grained readout of network criticality than the summary branching-ratio and Kappa metrics reported above. Pooled avalanche size and duration distributions (Figure 4A and B) overlapped closely between long-survival and early-death wells and were well described by heavy-tailed, approximately power-law distributions spanning three to four orders of magnitude in density, consistent with near-critical avalanche dynamics in both phenotypes while networks remained electrophysiologically active.

**Figure 4.**
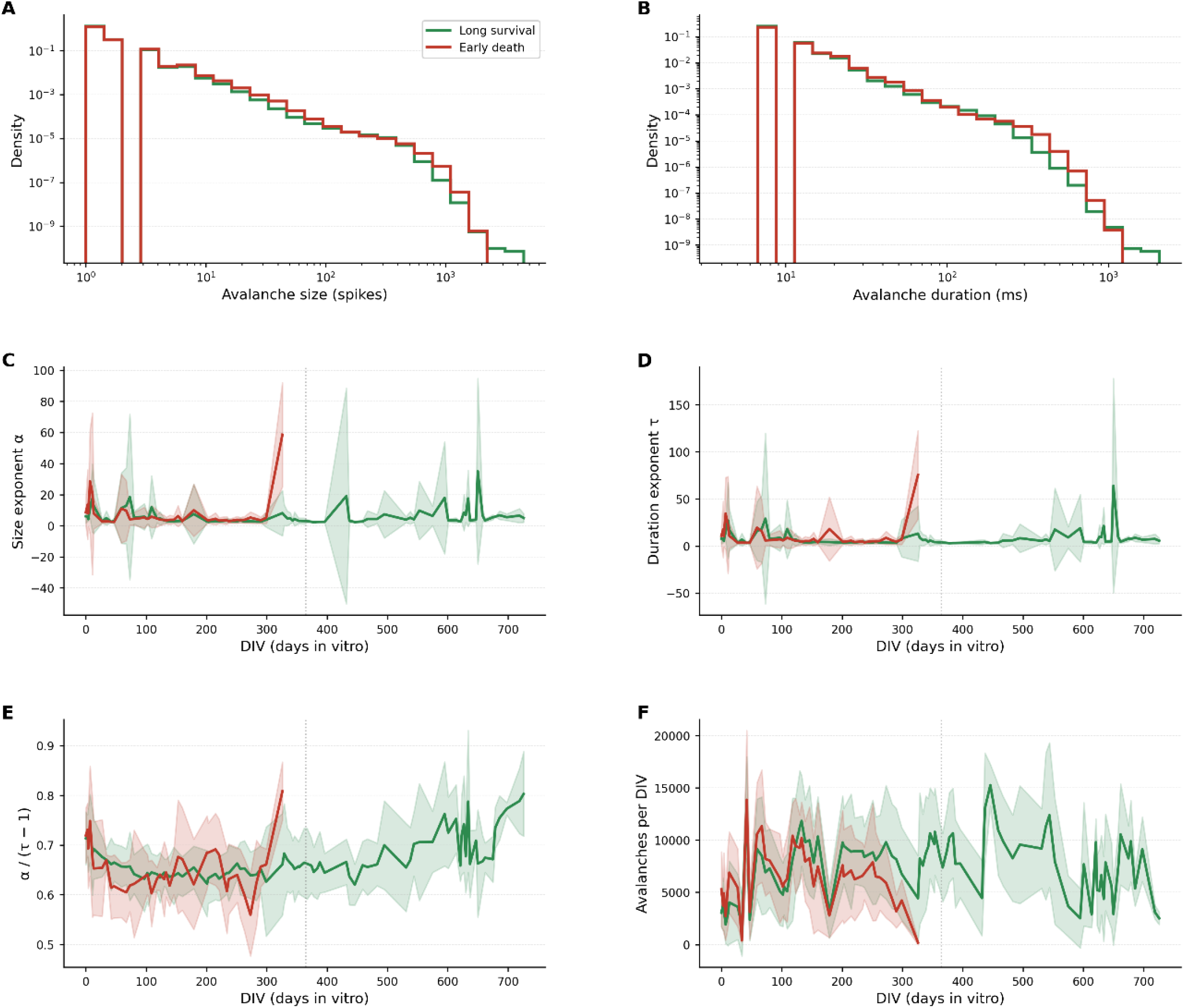
Neuronal avalanche statistics. (A) Probability density of avalanche size (spikes per avalanche), log–log axes, pooled across the full recording for long-survival (green) and early-death (red) wells; both groups show heavy-tailed, approximately power-law distributions spanning ∼4 orders of magnitude in density. (B) Probability density of avalanche duration (ms), as in (A). (C) Avalanche size exponent α, estimated within sliding DIV windows, as a function of DIV for each group (shaded band: between-well dispersion). (D) Avalanche duration exponent τ, as in (C). (E) Scaling relation α/(τ−1), the internal-consistency test for critical avalanche dynamics, as a function of DIV. (F) Avalanches detected per DIV. Vertical dotted line marks DIV 365.

As in the other longitudinal panels, each well’s contribution to Figure 4C**–F** ends at its own death or censoring DIV. Within this censored window, the size exponent α and duration exponent τ, estimated within sliding DIV windows (Figure 4C and D), were noisy throughout the recording in both groups, reflecting the comparatively small number of avalanches captured within each window; wide uncertainty bands appear intermittently in both long-survival and early-death wells, including during early development. The most prominent single feature in either exponent is a sharp excursion in early-death wells in the final weeks before their individual death DIV (approximately DIV 300–330), with α and τ transiently reaching values well above their typical range and markedly widened uncertainty bounds. Long-survival wells did not show a comparable systematic destabilization at any point up to DIV ∼730. This pattern was mirrored in the avalanche size–duration scaling relation α/(τ−1) (Figure 4E), a standard internal-consistency test of critical avalanche dynamics. Both groups tracked a similar, slowly evolving trajectory around 0.6–0.7 for most of their active recording, indicating that the exponents remained mutually consistent with scale-free avalanche propagation in both phenotypes. In early-death wells, this relation showed a comparatively larger, more erratic excursion in the weeks immediately preceding death; long-survival wells showed a more gradual upward drift over the second half of the recording, without a comparable destabilization.

The proximate driver of this instability is evident in the avalanche occurrence rate (Figure 4F): early-death wells showed a decline in avalanches detected per DIV over the course of the recording, with rates falling toward low values as wells approached their individual death DIV. Long-survival wells sustained higher and more variable avalanche rates (typically several thousand to over 10,000 per DIV) throughout most of the recording. Because reliable power-law exponent estimation requires a sufficient number of avalanche events, this decline in avalanche count in early-death wells plausibly underlies the instability of α, τ and their scaling relation in the period immediately before death. However, when tested as an independent functional survival associated-feature in the Cox proportional-hazards framework, avalanche occurrence rate at DIV 100–300 did not reach statistical significance after FDR correction (HR = 0.78, 95% CI 0.52–1.17, q = 0.339; **Table 1**; **Table S4**), indicating that, on its own, this metric should be regarded as a descriptive, exploratory signature of impending network failure rather than a validated functional survival associated-feature in this dataset.

### Multi-metric view of functional survival associated-features of network death

**Table 1** consolidates the individual functional survival associated-features of network death identified above, combining the FDR-corrected Cox proportional-hazards results (**Table S4**) with the Pearson-correlation analyses of Figures 1B **and 3I**. Seven features reached significance in the Cox framework, spanning burst structure, activity and criticality; three additional criticality metrics, the longitudinal slopes of branching ratio, CV of ISI and Kappa index, reached significance as simple correlates of survival time but were not among the thirteen features tested in the Cox model. Avalanche occurrence rate showed a clear descriptive separation between groups (Figure 4F) but did not reach significance in either framework.

Reported Pearson r values for burst, activity and criticality metrics are drawn from Figures 1B **and 3I**; Cox hazard ratios (HR, per 1 SD change in the z-scored feature) and FDR-corrected q-values are drawn from **Table S4**. HR > 1 indicates association with a higher hazard (earlier death); HR < 1, a lower hazard (later death). Only branching ratio (slope), CV of ISI (slope), Kappa index (slope) and burst duration (DIV 100–300) reached significance by simple correlation; seven further metrics reached significance in the Cox framework. Avalanche occurrence rate is included for completeness but did not reach significance in either analysis.

**Table 2** organizes these metrics by the recording window in which they become informative, outlining a staged view of a well’s trajectory toward death. Even within the earliest window (DIV 20–100), when conventional group-level trajectories still overlap substantially (Figure 2) and most simple correlations are weak (Figure 1B), the Cox model identifies early mean firing rate and synchrony as significant functional survival associated-features, indicating that modest, consistent differences are already present before they become visually apparent in group-averaged trajectories. The DIV 100–300 window carries the largest concentration of validated functional survival associated-features, including burst duration, inter-burst interval, synchrony, branching ratio and Kappa index and is where the branching-ratio and Kappa slopes computed across the full recording also begin to diverge most clearly between groups. The DIV >300 window, populated predominantly by long-survival wells together with the small number of early-death wells that persist long enough to contribute data there, functions as a confirmatory window in which branching ratio, Kappa index and DFA exponent differences become significant by simple comparison (Figure 3F**–H**), consistent with terminal network collapse rather than early prediction.

**Table 2.**
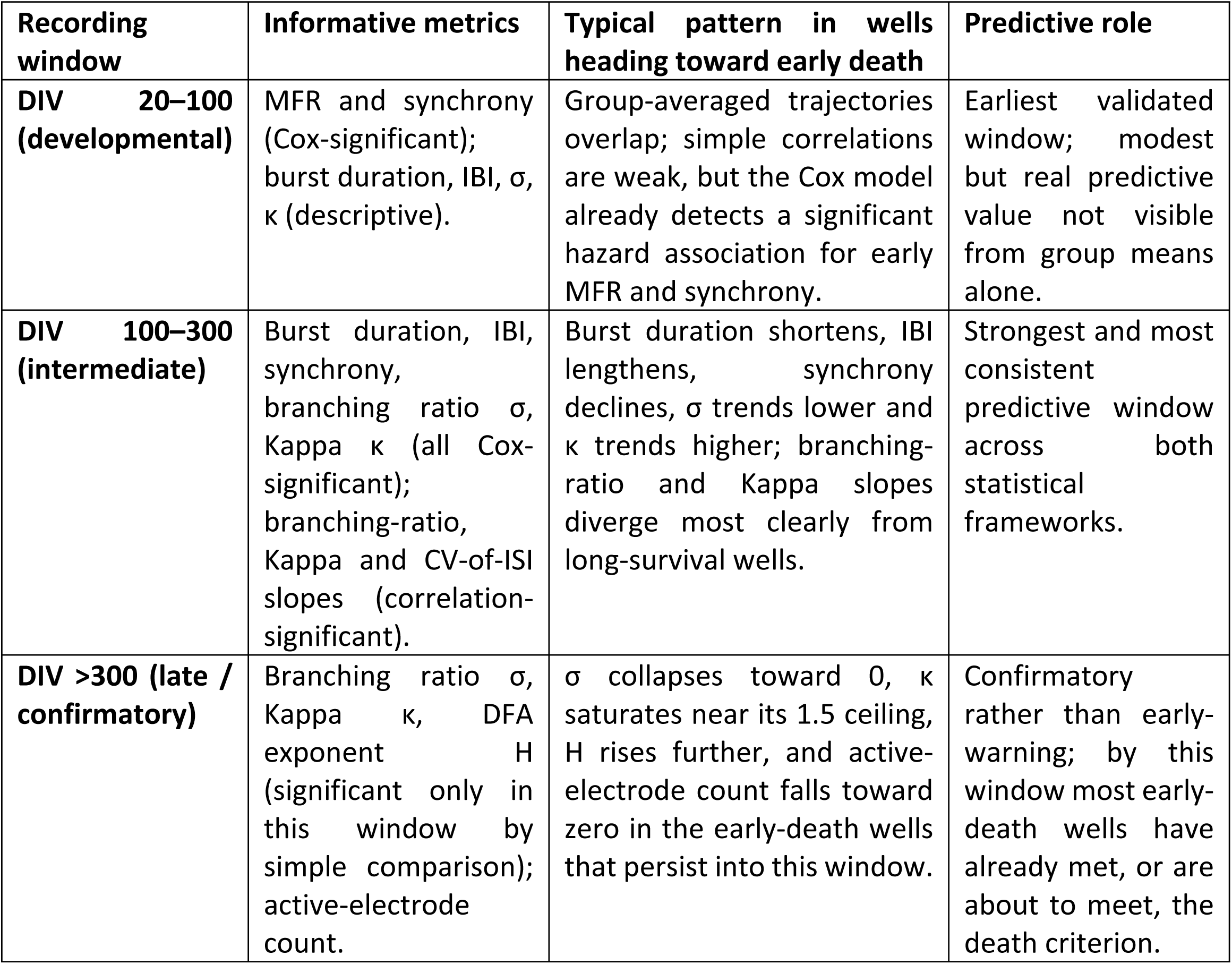
A staged view of predictors of network death, organized by recording window.

| Recording window | Informative metrics | Typical pattern in wells heading toward early death | Predictive role |
| --- | --- | --- | --- |
| **DIV 20–100 (developmental)** | MFR and synchrony (Cox-significant); burst duration, IBI, $\sigma$, $\kappa$ (descriptive). | Group-averaged trajectories overlap; simple correlations are weak, but the Cox model already detects a significant hazard association for early MFR and synchrony. | Earliest validated window; modest but real predictive value not visible from group means alone. |
| **DIV 100–300 (intermediate)** | Burst duration, IBI, synchrony, branching ratio $\sigma$, Kappa $\kappa$ (all Cox-significant); branching-ratio, Kappa and CV-of-ISI slopes (correlation-significant). | Burst duration shortens, IBI lengthens, synchrony declines, $\sigma$ trends lower and $\kappa$ trends higher; branching-ratio and Kappa slopes diverge most clearly from long-survival wells. | Strongest and most consistent predictive window across both statistical frameworks. |
| **DIV >300 (late / confirmatory)** | Branching ratio $\sigma$, Kappa $\kappa$, DFA exponent $H$ (significant only in this window by simple comparison); active-electrode count. | $\sigma$ collapses toward 0, $\kappa$ saturates near its 1.5 ceiling, $H$ rises further, and active-electrode count falls toward zero in the early-death wells that persist into this window. | Confirmatory rather than early-warning; by this window most early-death wells have already met, or are about to meet, the death criterion. |

## Discussion

This study provides a longitudinal view of the functional organization and decline of primary cortical networks over an unusually extended period. Previous studies have established the developmental sequence through which dissociated cortical cultures progress from sparse spiking to coordinated bursting and synchrony, predominantly during the first weeks in vitro,^4,6^and sealed culture systems have demonstrated that cortical networks can retain spontaneous electrical activity for more than one year.^12,13^ The present work extends these observations by characterizing multiple dimensions of network activity across individual wells for up to approximately 730 DIV, directly challenging the widespread assumption that such cultures remain electrophysiologically functional only for 4–8 weeks, an assumption that has shaped the design of virtually all MEA-based pharmacological and neurotoxicity studies to date.^6,22(p),23,24^ Long-survival wells maintained coherent burst structure and organized dynamics through DIV 500–600, with gradual decline thereafter, and some wells even showed renewed increases in activity at specific time points, suggesting ongoing plasticity rather than passive decay. The principal contribution of this work is therefore not simply the demonstration of prolonged activity, but the identification of electrophysiological features associated with contrasting trajectories of long-term functional maintenance.

A central implication of our findings is that firing rate alone provides an incomplete measure of long-term network integrity. Wells with different survival trajectories shared a broadly similar developmental phase and overlapping early firing-rate trajectories and the simple correlation between early MFR and survival time was weak (r ≈ −0.10 to −0.12). Only the Cox proportional-hazards model, which explicitly incorporates the timing of network death, revealed early MFR as a significant, independent predictor (HR = 1.36, q < 0.001), a modest but consistent early elevation in firing rate that was not visible from group averages alone. Burst organization separated the groups far more clearly: early-death wells showed longer inter-burst intervals from early DIVs despite comparable firing rates, and burst duration during DIV 100–300 was the strongest activity-related predictor by simple correlation (r ≈ 0.38), independently confirmed by Cox analysis (HR = 0.78, q < 0.001). This indicates that the richness of individual burst events is more informative about long-term viability than overall firing rate, consistent with prior MEA work showing that burst structure and coordination carry information about network state that average firing rate cannot capture.^25,26^

This dissociation between firing rate and burst structure has practical implications for MEA-based screening. An intervention that raises firing rate without improving burst duration or reducing inter-burst interval may not represent genuine network benefit,^25,27^ whereas moderate firing accompanied by stable bursting and synchrony may indicate more sustained network health.^28,29^ Current neurotoxicity assays based mainly on firing rate may therefore misclassify beneficial compounds, arguing for a broader, multidimensional functional panel, which incorporates firing, bursting, synchrony and electrode recruitment as a more informative standard MEA readout.^25^

Criticality-related measures provided the most mechanistically informative dimension of this profile, supported both by window-averaged Cox modelling and by the longitudinal trajectory of each metric.^30^ The branching ratio σ, reflecting how spiking activity propagates through the network, was the strongest Cox-validated criticality predictor at DIV 100–300 (HR = 0.34, q < 0.001) and its slope across the full recording was the single strongest correlate of survival time among all metrics tested (r ≈ 0.48). The Kappa index was the strongest single Cox predictor overall (HR = 3.02, q < 0.001), with early-death wells drifting toward κ ≈ 1.5 particularly as they approached death, while long-survival wells remained closer to the κ = 1 critical line for most of the recording. This is consistent with the broader hypothesis that critical-like dynamics support efficient information transmission and dynamic range,^30–32^ and the observed supercritical drift reflects a loss of structured avalanche dynamics analogous to the loss of critical dynamics that precedes functional collapse in models of neurodegeneration. The coefficient of variation of the inter-spike interval added further evidence: although not significant as a window-averaged predictor, its slope was the second-strongest correlate of survival overall (r ≈ 0.46), reinforcing that trajectory, not just instantaneous level, carries prognostic value. Group-level differences in σ and κ reached significance well before conventional metrics like MFR or burst rate, offering a prospective window of months into network fate; the DFA Hurst exponent, by contrast, only differed significantly between groups after DIV 300 (p = 0.031), making it more useful as a late-stage confirmatory measure than an early warning sign.

The avalanche-level analysis reinforced this picture. Avalanche size and duration statistics, and their scaling relation, were closely matched between phenotypes while networks remained active, indicating that both organize activity into scale-free cascades for most of their functional lifespan; the clearest departure was a transient destabilization of these exponents in early-death wells in the weeks immediately preceding death, alongside a decline in avalanche occurrence rate. This occurrence rate, however, did not reach significance as an independent Cox predictor (HR = 0.78, q = 0.339), suggesting either a genuinely weaker association than the propagation-based metrics or reduced power given the limited number of events, at present it should be regarded as a descriptive signature tracking the same underlying decline captured by σ and κ, rather than a validated independent predictor. More broadly, heavy-tailed distributions are compatible with scale-free organization but do not by themselves establish a power law or criticality without formal goodness-of-fit testing, comparison with alternative distributions, and complementary scaling analyses,^33(p),34^ since power-law-like distributions can also emerge in non-critical systems and branching estimates are affected by subsampling.^35^ The observed changes are therefore best described as alterations in criticality-related dynamics rather than definitive transitions between critical and non-critical states, and declining avalanche counts near functional silencing both signal deterioration and reduce the precision of exponent estimation.

The spontaneous segregation of wells into two reproducible survival phenotypes positions long-term cortical MEA cultures as a tractable model for neuronal ageing research,^36–38^ distinct from biochemical ageing models based on senescence markers or transcriptomic profiles,^39–41^ since MEA recordings provide continuous, non-destructive functional monitoring without perturbing the cultures. The metrics identified here, particularly σ and κ, could serve as objective endpoints for testing interventions designed to extend network lifespan or preserve critical-state organization. For pharmacological screening, the DIV 100–300 predictive window means a compound’s effect on network fate could be detected within 3–6 months rather than over the full two-year recording; because σ and κ carry mechanistic information about network state rather than activity level alone, they are better positioned to distinguish neuroprotective from simply stimulatory effects,^30^ and the multi-well format allows treatment and vehicle-control wells to be maintained indefinitely for within-plate comparison.

Taken together, Cox proportional-hazards modelling with FDR correction identified Kappa index and branching ratio at DIV 100–300, together with burst duration and inter-burst interval at the same window, and synchrony and mean firing rate at DIV 20–100, as independent functional predictors of survival time. This combination of conventional activity measures with criticality-derived metrics, validated through both simple correlation and formal survival modelling, establishes a quantitative framework for identifying functional biomarkers of network decline that outperform firing rate alone, and positions long-term cortical MEA cultures as a tractable platform for neuronal ageing research and for electrophysiology-guided screening of neuroprotective and anti-ageing compounds.

## Materials and methods

### Primary Cortical Neuron Culture

Primary cortical (CTX) neurons were dissected from 16-day-old embryos (E16) of Wistar rats (Charles River Laboratory). Briefly, the brains were extracted and kept in Leibovitz’s L-15 medium (Gibco) on ice. The cortices were separated and transferred to Neurobasal Plus medium (Gibco). Cell dissociation was performed mechanically using a crystal Pasteur pipette, and the resulting suspension was centrifuged for 5 minutes at 500 × g. The supernatant was aspirated and the pellet was resuspended in culture medium.

Culture medium was prepared by supplementing Neurobasal Plus (Gibco) with 1× GlutaMAX (Gibco), 1× B-27 Plus supplement (Gibco), and 1% penicillin/streptomycin (Gibco). All substrates and MEA wells underwent a standardized three-step coating protocol. Step 1: incubation with 0.1% polyethylenimine (PEI, Mw ∼25,000; Sigma) in 1× borate buffer for 1 hour, followed by three washes with sterile H₂O (Gibco). Step 2: incubation with 1 mg/mL poly-D-lysine (Gibco) in Dulbecco’s phosphate-buffered saline (DPBS, Gibco) for 1 hour, followed by three washes with sterile H₂O and air-drying. Step 3: incubation with 1% Geltrex (Gibco) in culture medium for 24 hours; Geltrex was removed immediately prior to cell seeding. Cells were seeded at a density of 10⁵ cells/well and maintained in a humidified incubator at 37 °C with 5% CO₂. Half-medium changes were performed at 4 DIV and subsequently at regular intervals (2 to 3 times per week).

### Ethics statement

The study used brain tissue collected in 2020 from E16 rat fetuses for the establishment of primary neuronal cultures. Pregnant rats that had undergone no previous experimental intervention were deeply anesthetized with isoflurane and subsequently euthanized by decapitation solely for fetal tissue collection. Immediately thereafter, E16 fetuses were collected, and their brains were dissected for the preparation of primary neuronal cultures. The fetuses were not maintained alive or subjected to any independent in vivo intervention. Under Article 3 of Directive 2010/63/EU and the corresponding Spanish legislation applicable at the time, Royal Decree 53/2013, the killing of animals solely for the use of their organs or tissues did not constitute an experimental procedure requiring project authorization. Therefore, no specific animal experimental project authorization was required for this tissue collection. The animals were obtained from an authorized breeder (Charles River Laboratories), and all procedures related to their housing, handling, and euthanasia were performed by competent personnel in accordance with the institutional and national animal-welfare requirements applicable in 2020.

### Multi-Electrode Array Recordings

Electrophysiological activity was recorded using a 24-well microelectrode array (MEA) plate with 16 electrodes per well (384 channels total), sampled at 12,500 Hz. Cells were seeded on top of the gold (Au) electrode surfaces at a density of 10⁵ cells/well (**Figure S1**). After DIV 15–25, spontaneous electrical activity was recorded for 5 minutes per session. Prior to each recording, the MEA plate was placed in the MEA reader and allowed to equilibrate thermally at 37 °C with 5% CO₂ for 15 minutes. Two independent culture preparations were tracked longitudinally over the entire culture lifetime. Recordings were performed at multiple time points spanning from early DIV through approximately DIV 700.

### Computational analysis

Raw voltage traces were stored as HDF5-formatted .mat files and processed with a custom Python pipeline using NumPy, SciPy, pandas, h5py, and matplotlib.

To ensure only wells with genuine network activity were included in the analysis, a two-step quality filter was applied. First, a well was considered active if, in every recording session between DIV 20 and DIV 40, at least 8 out of 16 electrodes exceeded a firing rate of 1 Hz. Wells failing this criterion were excluded entirely. Second, a well was declared dead at the first DIV from which five consecutive sessions showed three or fewer active electrodes. Active wells were then classified into two groups; a) early death, where well death occurred before DIV 365 (within the first year in vitro) and b) long survival, where well death occurred at DIV 365 or later, or the well remained active at the final recording session.

For each active well, the following parameters were computed at every recorded DIV and tracked longitudinally: mean firing rate (MFR), calculated as the mean firing rate across active electrodes (FR > 1 Hz) at each time point; burst rate, burst duration, and inter-burst interval (IBI), derived from the population-averaged signal; network synchrony, quantified as the mean pairwise Pearson correlation across electrodes within a well; and criticality metrics, including the coefficient of variation of the inter-spike interval (CV_ISI), Fano factor, branching ratio (σ), kappa index (κ) and the DFA Hurst exponent (H). All metrics were analyzed across the full recording period (DIV > 18), compared between the two survival groups, and aligned to the individual death DIV of each well to identify patterns predictive of network decline.

### Electrophysiological and criticality metrics

To comprehensively characterize the temporal evolution of neuronal network activity and identify electrophysiological signatures associated with long-term survival, a set of conventional activity descriptors and criticality-related metrics was calculated for each recording. Conventional metrics quantified firing activity, bursting behavior, and network synchrony, whereas criticality metrics evaluated the dynamical state of the network and its proximity to critical behavior. All analyses were performed independently for each well at every recording time point.

### Mean Firing Rate (*MFR*)

*MFR* (**Eq. 1**) is the most widely used descriptor of neuronal activity in MEA studies and provides a global estimate of the excitability of the neuronal network. Higher *MFR* values generally indicate increased spontaneous neuronal activity, although elevated firing alone does not necessarily imply improved network organization or functional stability. The *MFR* was calculated as the average firing rate (*FR*, **Eq. 2**), that measures the activity of individual electrodes and reflects the local excitability of the neuronal population of all active electrodes (*i*) within each well (electrodes exhibiting *FR* > 1 Hz).

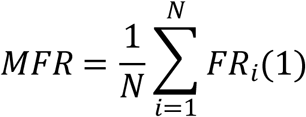

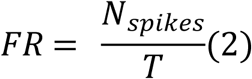

### Burst rate (*BR*) and duration (*BD*)

*BR* (**Eq. 3**) quantifies the average frequency of local bursting events across active electrodes and is commonly used as an indicator of functional maturation and connectivity and longer *BD*s (**Eq. 4**) generally indicate that neuronal assemblies can sustain coordinated activity for extended periods, reflecting stronger functional connectivity.

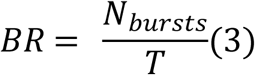

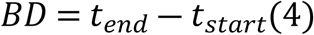

### Inter-burst interval (*IBI*)

*IBI* (**Eq. 5**) reflects the temporal spacing between synchronized events. Shorter IBIs indicate more frequent coordinated activity, whereas prolonged IBIs suggest reduced network excitability or impaired synchronization.

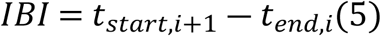

### Network synchrony

Higher synchrony index (*SI*, **Eq. 6**) values indicate stronger temporal coordination across the neuronal population and greater functional connectivity. It was quantified as the mean pairwise Pearson correlation coefficient (*r_ij_*, **Eq. 7**) between all active electrode spike trains.

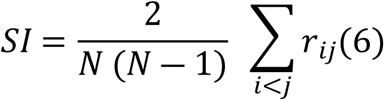

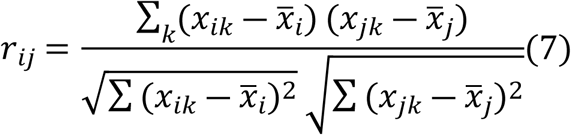

### Coefficient of variation of inter-spike interval (*CV_ISI_*)

*CV_ISI_* (**Eq. 8**) measures spike train irregularity, as a fraction of the standard deviation (*σ_ISI_*)and the mean (µ*_ISI_*). A value close to 1 is expected for a Poisson process, values below 1 indicate regular firing, and values above 1 reflect increasingly irregular or burst-like firing.

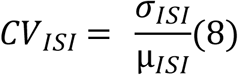

### Fano factor (*FF*)

It is used to quantify the spike-count variability (**Eq. 9**). A Poisson process has FF= and values greater than one indicate increased temporal variability, whereas values below one indicates more regular firing.

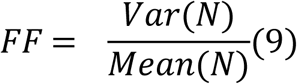

### Branching ratio

The branching ratio (*σ*, **Eq. 10**) quantifies how efficiently activity propagates through the neuronal network, as *A_t_* is the number of active units at time *t* and *A_t_*_+1_ is the activity in the subsequent time bin. Networks operating near the critical state exhibit *σ* ≈ 1. Values below one indicates subcritical dynamics, whereas values above one indicates supercritical propagation.

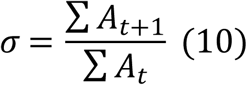

### Avalanche size and duration distribution

Neuronal avalanche size (*S*, **Eq. 11**) was defined as the total number of spikes within each avalanche, being *τ* the avalanche size exponent. Power-law scaling indicates scale-free organization, which is a hallmark of critical neuronal dynamics.

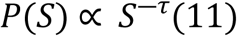

The avalanche duration exponent (*T*, **Eq. 12**) provides complementary information about the temporal organization of neuronal activity, with *α* the temporal scaling exponent.

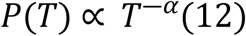

### Kappa index

The Kappa index (*κ*, **Eq. 13**) was calculated by comparing the empirical cumulative avalanche distribution with the theoretical power-law prediction, where *F_empirical_* is the cumulative empirical distribution, *F_t_*_ℎ*eory*_ is the expected cumulative power-law distribution and *m* is the number of evaluation points. A value of *κ* ≈ 1 indicates dynamics close to criticality. Values below one corresponds to subcritical dynamics, whereas values above one indicates increasingly supercritical network behavior.

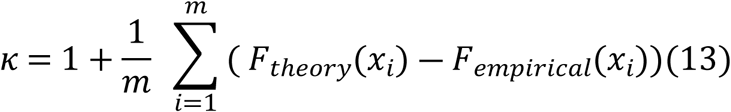

### Detrended fluctuation analysis (DFA) and Hurst exponent (*H*)

Long-range temporal correlations were quantified using detrended fluctuation analysis, where *F*(*n*) is the root-mean-square fluctuation after detrending, *n* is the window length and *H* is the Hurst exponent.

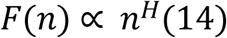

DFA is robust against non-stationary signals and is widely used to characterize scale-free temporal organization in neuronal activity. The *H* obtained from DFA characterizes long-range temporal dependence. An exponent of *H* = 0.5 indicates an uncorrelated signal consistent with white noise, whereas *H* < 0.5 reflects anti-persistent temporal dynamics. Values of *H* > 0.5 indicate persistent long-range temporal correlations, with exponents approaching *H* ≈ 1 representing strong temporal organization that is characteristic of neuronal systems operating near a critical state. Because persistent long-range temporal correlations are considered a hallmark of healthy neuronal network dynamics, the DFA exponent is widely used as an indicator of proximity to criticality in neural systems.

### Statistical analysis of predictors of network death

To identify individual electrophysiological and criticality-derived predictors of network death, and to test whether combining predictors improved on any single metric, the per-well feature tables described above (metric values averaged within the DIV 20–100 and DIV 100–300 windows) were analyzed in Python 3.9 using lifelines (v0.30) for survival modeling, scikit-learn (v1.2) for the combined classifier and an in-house implementation of the Benjamini-Hochberg procedure for multiple-comparison correction.

For each of 13 candidate features, a univariate Cox proportional-hazards model was fit relating the z-scored feature value to the instantaneous hazard of a well reaching the death criterion. To control the false discovery rate across the family of 13 univariate tests, raw p-values were adjusted using the Benjamini-Hochberg procedure; features with an adjusted q-value < 0.05 were considered statistically significant.

To test whether combining multiple criticality-derived features improved prediction of network fate beyond any single metric, an L2-regularized (ridge) logistic regression was fit to classify wells as early-death versus long-survival, using five DIV 100–300 features selected for individual predictive strength and conceptual non-redundancy: branching ratio (σ), Kappa index (κ), burst duration, inter-burst interval, and avalanche occurrence rate. All features were z-scored prior to fitting.

## Acknowledgements

This work was supported by grant PID2022-137363OA-I00 funded by MCIN/AEI/10.13039/501100011033. A.E. was supported by a postdoctoral fellowship from the Provincial Council of Gipuzkoa under the Fellows Gipuzkoa Program. We acknowledge financial support from IKUR Strategy (NanoNeuro) under a collaboration agreement between Ikerbasque Foundation and the Materials Physics Center, and BioGipuzkoa, respectively, on behalf of the Department of Science, Universities and Innovation of the Basque Government. Funding supporting F.J.G.-B. was provided by the Basque Government (grants #2024333026, and #2025333026), Bioef (#BIO17/ND/023-BD), and IKERBASQUE Basque Foundation for Science (#PP/2022/003). L.R.G. was supported by a predoctoral fellowship from the Basque Government (#PRE_2021_1_0125). The authors thank Marek Grzelczak for discussion.

Keywords: Multi-electrode arrays, Neuronal criticality, Cortical network development, Survival analysis, Neuronal ageing

## CRediT authorship contribution statement

**Ane Escobar:** Conceptualization, Methodology, Supervision, Investigation, Formal analysis, Funding acquisi-tion, Project administration, Data curation, Writing – original draft, Writing – review & editing. **Laura Rodríguez-Gómez:** Methodology, Writing – review & editing. **Francisco Javier Gil-Bea:** Conceptualization, Supervision, Validation, Funding acquisition, Writing – review & editing.

## Data Availability Statement

The data that support the findings of this study are available from the corresponding author upon request.

## Supporting information captions

**Figure S1.** Representative micrographs of primary cortical neurons cultured on microelectrode array (MEA) electrodes. Representative bright-field images acquired at different magnifications showing primary cortical neurons growing directly on the gold (Au) recording electrodes of the 24-well MEA plates. Scale-bar = 100 µm.

**Figure S2.** Flowchart of well inclusion and survival classification. Wells were included only if at least 8 of 16 electrodes were active (MFR > 1 Hz) during every recording between DIV 20 and DIV 40. Well death was defined as the first DIV followed by five consecutive recordings with ≤3 active electrodes. Included wells were classified as early death (death before DIV 365) or long survival (death at or beyond DIV 365, or surviving until the end of the experiment).

**Table S1.** Summary of experimental design and well classification.

**Table S2.** Electrophysiological parameters analyzed.

**Table S3.** Criticality metrics analyzed.

**Table S4.** Univariate Cox proportional-hazards predictors of network death, ranked by false discovery rate (FDR)-corrected q-value (Benjamini-Hochberg, 13 tests). Hazard ratios (HR) are per 1 SD change in the z-scored feature; HR > 1 indicates association with a higher hazard (earlier death), HR < 1 with a lower hazard (later death). Standard errors are clustered by culture. Sample size (n = 31–32 per feature), raw p-values, and concordance indices are reported in the main text (Table 1).

